# Are fleas highly modified Mecoptera? Phylogenomic resolution of Antliophora (Insecta: Holometabola)

**DOI:** 10.1101/2020.11.19.390666

**Authors:** Karen Meusemann, Michelle Trautwein, Frank Friedrich, Rolf G. Beutel, Brian M. Wiegmann, Alexander Donath, Lars Podsiadlowski, Malte Petersen, Oliver Niehuis, Christoph Mayer, Keith M. Bayless, Seunggwan Shin, Shanlin Liu, Ondrej Hlinka, Bui Quang Minh, Alexey Kozlov, Benoit Morel, Ralph S. Peters, Daniela Bartel, Simon Grove, Xin Zhou, Bernhard Misof, David K. Yeates

## Abstract

Insect orders have been defined and stable for decades, with few notable exceptions (*e.g*., Blattodea and Psocoptera). One of the few remaining questions of order-level monophyly is that of Mecoptera in respect to the phylogenetic placement of Siphonaptera (fleas). We used a large set of transcriptomic nucleotide sequence data representing 56 species and more than 3,000 single-copy genes to resolve the evolutionary history of Antliophora, including fleas (Siphonaptera), scorpionflies and relatives (Mecoptera), and true flies (Diptera). We find that fleas and mecopterans together are the sister group of flies. However, our data and/or analyses are unable to distinguish whether fleas are sister to a monophyletic Mecoptera, or whether they arose from within extant mecopteran families, rendering Mecoptera paraphyletic. We did not detect parameter bias in our dataset after applying a broad range of detection methods. Counter to a previous hypothesis that placed fleas within Mecoptera as the sister group to wingless boreids (snow fleas), we found a potential sister group relationship between fleas and the enigmatic family Nannochoristidae. Although we lack conclusive evidence, it seems possible that fleas represent the most-species rich group of modern mecopterans and that their parasitic lifestyle and morphological adaptations have simply made them unrecognizable in respect to their order-level classification.

## Introduction

The evolutionary relationships of insects are increasingly well-known as the result of large-scale phylogenomic analyses (Misof et al., 2014; Peters et al., 2017; Johnson et al., 2018; Wipfler et al., 2019; Evangelista et al., 2019; Kawahara et al., 2019; McKenna et al., 2019; Vasilikopoulos et al., 2020). The phylogenetic relationships within Holometabola, insects that undergo complete metamorphosis, are particularly well-understood and have been consistently confirmed by morphological and molecular work over the last decade (Wiegmann et al., 2009; McKenna & Farrell, 2010; Beutel et al., 2011; Trautwein et al., 2012; Peters et al., 2014). An exception to the generally robustly resolved phylogenetic relationships within Holometabola are the inconsistent phylogenetic relationships in Antliophora: a group that comprises flies (Diptera), fleas (Siphonaptera) and the relatively less well known scorpionflies and relatives (Mecoptera) (Whiting 2002; Wiegmann et al., 2011; Misof et al., 2014). The phylogeny of Antliophora is a compelling area of focus because it includes one of the last remaining issues of order-level monophyly within Insecta: the putative paraphyly of Mecoptera due to the inclusion of fleas.

Within Antliophora, flies (Diptera) represent the largest order–including more than 158,000 described extant species. In fact, Diptera is one of the four largest orders of insects overall, making up ~ 10% of known species diversity on Earth. The closest relatives of flies have traditionally been considered either fleas (Siphonaptera) (Boudreaux 1979), scorpionflies (Mecoptera, Hennig 1969), or fleas and scorpionflies together (Ross 1965; Kristensen 1991). Fleas are wingless, blood-sucking, obligate parasites, primarily of mammals (although a small number of ~ 2,500 total species parasitize birds, see Zhu et al., 2015). Mecopterans are a small heterogeneous order, comprised of only ~ 600 described extant species (Whiting 2002), including large, showy scorpionflies (Panorpidae, Figure 1), as well as less conspicuous groups such as wingless, ice-dwelling snow scorpionflies (*i.e.,* snow fleas, Boreidae, Figure 1).

**Figure 1.**
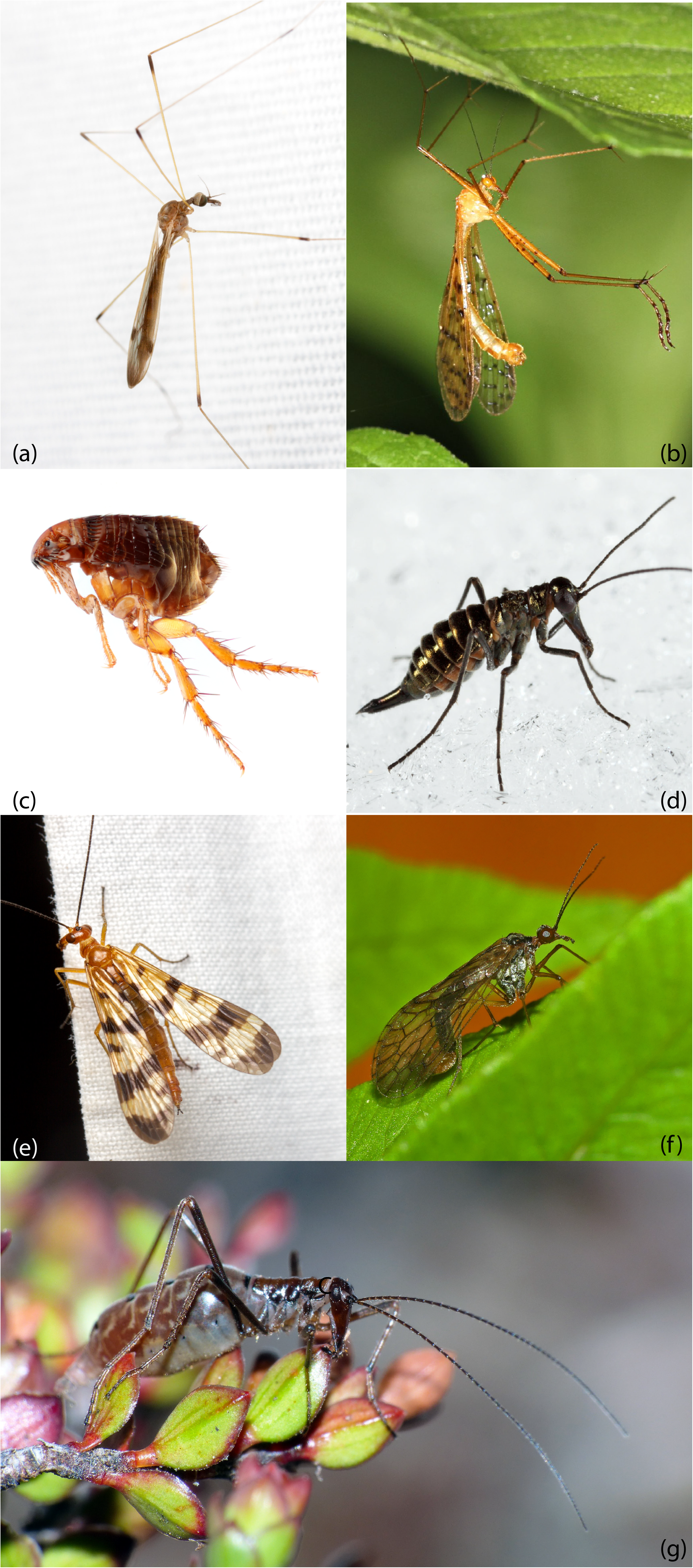
Photos of antliophoran representatives. **(a)** A crane fly (Diptera: Tipulidae: *Helius flavipes*) and **(b)**. a hanging fly (Mecoptera: Bittacidae: *Bittacus pilicornis*) exhibit superficial similarity. Photos by Matthew Bertone. **(c)** A flea (Siphonaptera) and **(d)** a snow scorpionfly or snow flea (Mecoptera: Boreidae: *Boreus brumalis)* were previously proposed as sister groups (Whiting, 2002). Photos by Matthew Bertone and Tom Murray, respectively. **(e)** A scorpionfly (Mecoptera: Panorpidae: *Panorpa* sp.) Photo by Matthew Bertone. **(f)** Nannochoristidae (Mecoptera: *Nannochorista dipteroides*) and **(g)** A wingless scorpionfly (Mecoptera: *Apteropanorpa tasmanica*) Photos by Simon Grove.

Mecopterans have been common and diverse since the early Permian until the Cretaceous, and their diversity has been reduced to the modern depauperate and relict fauna since the Paleocene (Novokshonov 2002; Grimaldi & Engel, 2005; Misof et al., 2014). The anatomical diversity of their fossil forms is great, from mosquito-like, two-winged species with elongate mouthparts (*Pseudopolycentropus*), to bizarre, wingless ectoparasites with very long (*Saurophthirus)* or very swollen (*Strashila*) legs. This fossil diversity is almost matched by extant forms, including variously disparate lineages, such as families with wingless cold-adapted species (Boreidae and Apteropanoridae), or families with large, flattened species resembling cockroaches with elongate male genitalic claspers (Meropeidae). Mecopterans also include groups that share general appearances with early diverging true flies: the hanging scorpionflies (Bittacidae) that resemble crane flies (Tipulidae) (Figure 1) and the enigmatic Nannochoristidae, whose aquatic larvae show similarities to fly and flea larvae (Figure 1) (*e.g*., Beutel & Friedrich, 2019).

Despite their anatomical diversity, Mecoptera are thought to be among the most morphologically generalized holometabolous insects, with three ocelli, mouthparts with biting mandibles, two pairs of unmodified palps, filiform multisegmented antennae, little differentiation of thoracic segments or wings, five tarsomeres, an eleven-segmented abdomen (Grimaldi & Engel, 2005), and larvae with compound eyes, which is unique in Holometabola, but widespread in nymphs of hemimetabolous insects. It is clear that the three modern orders Diptera, Siphonaptera, and Mecoptera share a common ancestor among a diverse assemblage of “mecopteroid” stem group antliophorans in the late Palaeozoic (Willmann 1987; Grimaldi & Engel, 2005; Misof et al., 2014).

A clear understanding of mecopteran relationships is crucial for understanding the early evolution of Antliophora. While a series of unambiguous synapomorphies establish the monophyly of Diptera and Siphonaptera, the monophyly of Mecoptera has been a subject of debate for decades, with controversy surrounding two families, Boreidae and Nannochoristidae (Willmann 1987; Whiting 2002; Beutel & Friedrich, 2019). During the 20th century, separate order status was conferred on both families, Neomecoptera (Hinton 1958) and Nannomecoptera (Hinton 1981), respectively; however, this was based on their unique morphological differences rather than a definitive phylogenetic placement separate from other scorpionflies. Aside from Boreidae and Nannochoristidae, the remaining mecopteran families are united in a group termed Pistillifera (Willmann 1987).

Nannochoristidae (Beutel & Friedrich, 2019) are unusual mecopterans with elongate, very slender aquatic larvae with a very smooth body surface (Figure 1). While superficially resembling other Mecoptera, the adults have some significant differences including the placement of the female genital opening, and the lack of the advanced pistilliferan sperm pump that is a synapomorphy of the other Mecopteran families (excluding Boreidae). This type of pump is not homologous to that occurring in Diptera (Hünefeld & Beutel, 2005), but nannochoristid sperm pumps do show some structural affinities with that of Siphonaptera (Mickoleit 2009). Nannochoristidae have even been suggested to be the lone sister group to Diptera (Wood & Borkent, 1989), though this idea has been refuted (*e.g*., Beutel & Baum, 2008; Beutel & Friedrich, 2019). Morphological evidence is presently equivocal regarding the placement of Nannochoristidae (Beutel & Friedrich, 2019). Early molecular work based on a few ribosomal, mitochondrial, and nuclear genes also recovered Nannochoristidae as sister to all remaining Mecoptera including Siphonaptera as sister to Boreidae (Whiting 2002).

The phylogenetic affinities of the family Boreidae have confounded previous efforts aiming at inferring the phylogenetic relationships of Mecoptera. Studies by Whiting and colleagues (Whiting et al., 1997; Whiting et al., 2002) placed the family as sister to fleas, rendering Mecoptera paraphyletic. Aside from the general similarities between fleas and boreids, such as winglessness and the ability to jump, they also share characters such as multiple sex chromosomes, ovaries lacking nurse cells (secondarily panoistic), and a propensity to feign death after jumping (Penny 1977; Kristensen 1999; Grimaldi & Engel, 2005). Despite these shared traits, recent results of morphological analyses did not provide evidence that suggest a close relationship between fleas and boreids (Beutel et al., 2008; Fabian et al., 2015). Additional evidence refuting this hypothesis was recovered in a molecular phylogeny by Wiegmann and colleagues (2009) who studied six nuclear genes and found monophyletic Mecoptera with Bittacidae as sister to the remaining Mecoptera. A large-scale transcriptome nucleotide sequence data-based phylogeny of insects (Misof et al., 2014) also recovered monophyletic Mecoptera with Boreidae as sister to the remaining families based on the phylogeneic analysis of more than a thousand single-copy protein-coding genes. Yet, supplementary results in the study by Misof and colleagues (2014) pointed to conflicting signal within the dataset, providing support for mecopteran paraphyly, with Nannochoristidae as sister group to Siphonaptera. In this case, an influence of model violation due to among-lineage heterogeneity and/or non-randomly distributed data within the dataset could not be excluded. Therefore, with the largest phylogenomic dataset yet applied to insect order-level phylogeny, the monophyly of Mecoptera could not be inferred conclusively.

The unresolved phylogenetic relationships of Mecoptera represent one of the very few remaining open questions of order-level monophyly within hexapods and are critical to understand the broader evolution of Antliophora. Phylogenetic ambiguity in Antliophora is caused by the varying support for multiple hypotheses of mecopteran phylogenetic relationships inferred using different data sources, and compounded by eroded phylogenetic signal over the long evolutionary history. Phylogenetic analyses of morphological data and of molecular datasets with few mitochondrial, ribosomal, or protein-coding nuclear genes and of large-scale transcriptome data implied different evolutionary scenarios for Antliophora. Here, we present results from analyzing up to 3,145 nuclear single-copy protein-coding genes extracted from transcriptomes and from draft genome assemblies from 56 species to resolve the phylogeny of Antliophora and to assess the monophyly of Mecoptera (Table 1). Our phylogenetic analyses include transcriptome nucleotide sequence data from twelve mecopterans, four siphonapterans, and thirty-two dipterans. Eight species from Lepidoptera and Trichoptera were included as the outgroup taxa for rooting of the inferred phylogenies (see Supplemetary Materials, Supplementary Tables S1–S3).

**Table 1.**
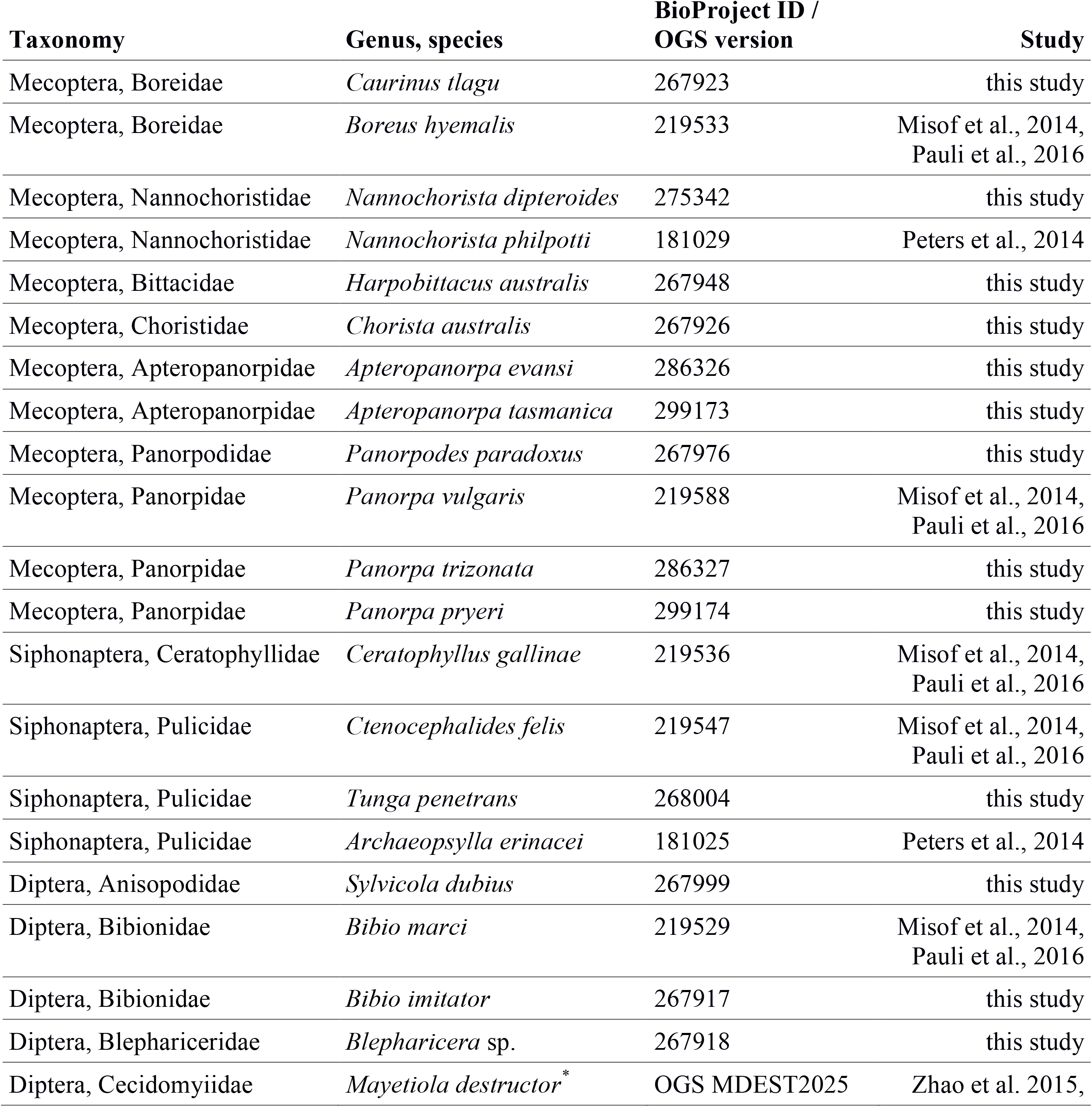
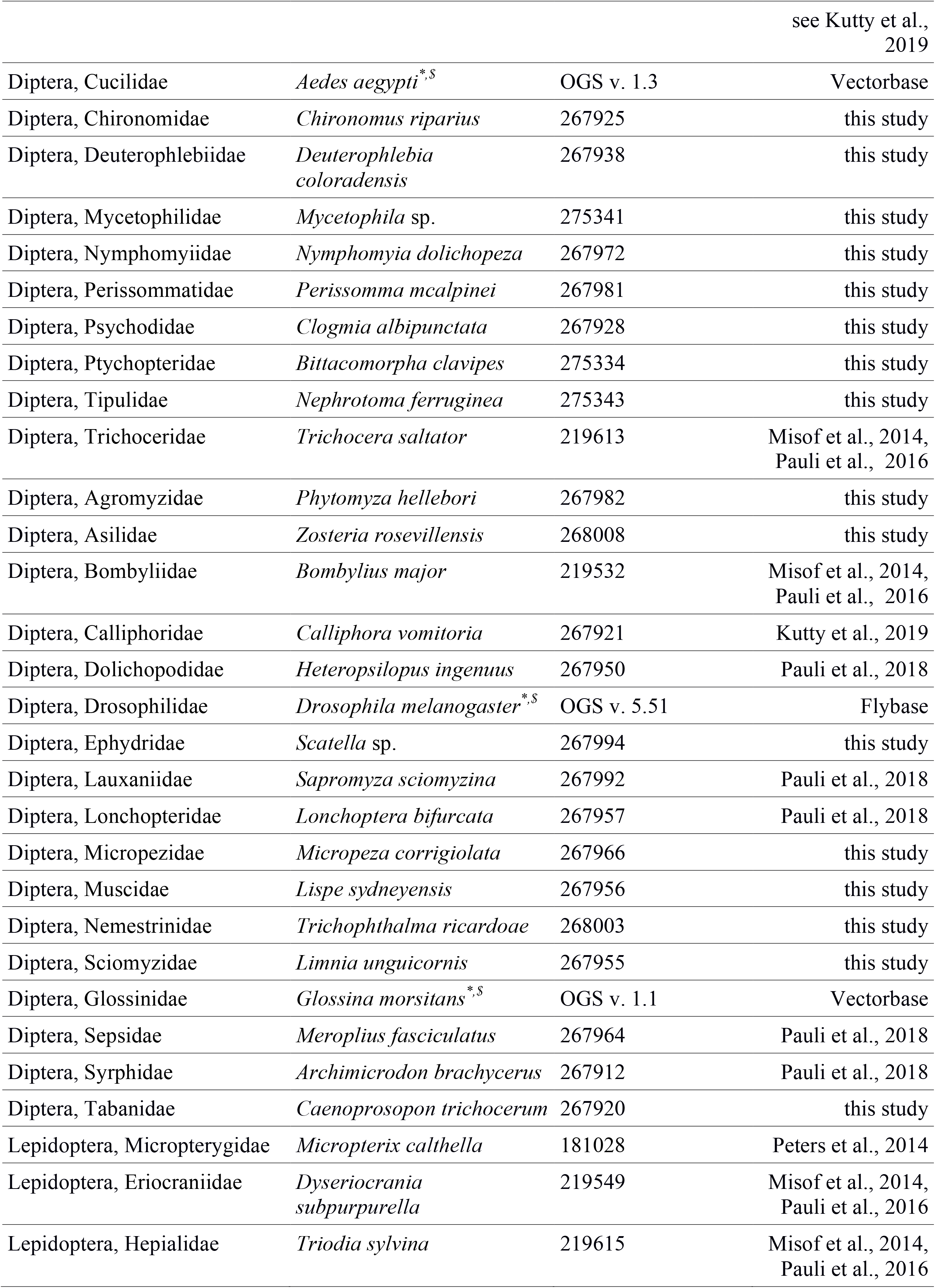
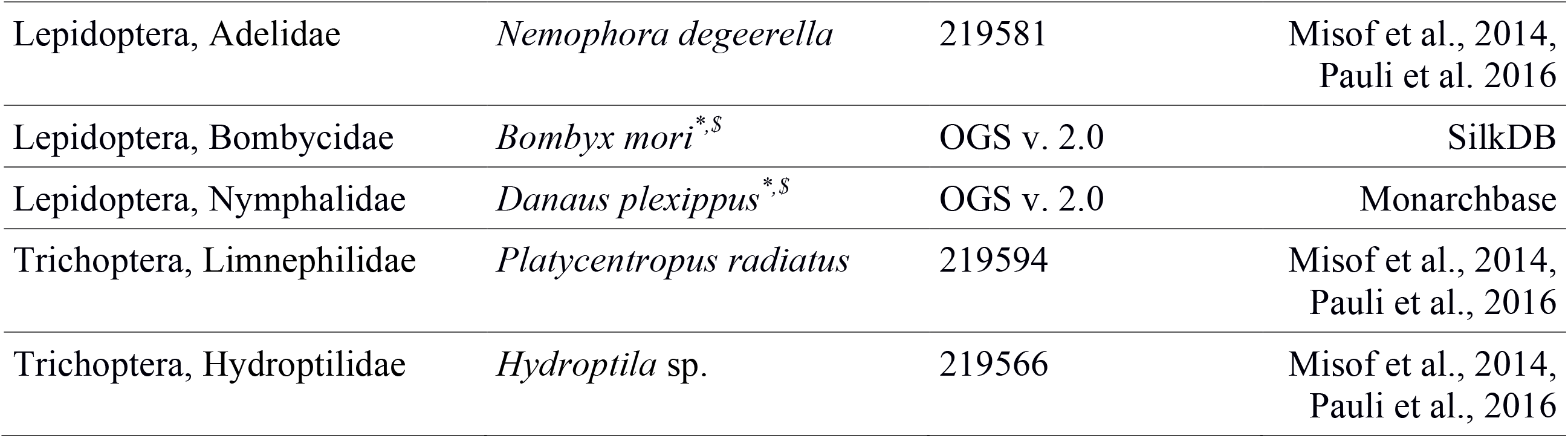
Species included in this study in our final analyses. BioProject IDs refer to the NCBI BioProject database (see Umbrella project “The 1KITE project: Evolution of insects”). For references, please refer to the main text and the Supplementary Materials. Details on collecting information, data sources, SRA and TSA accession numbers excluded species during analyses can be found in the Supplementary Materials: Supplementary Table S1–S4. *official gene set (OGS) version from available genomes, ^$^ species included as reference species in the ortholog set, see Pauli et al. (2018).

## Methods Summary

We collected specimens and generated new RNASeq data from nine species of Mecoptera, one species of Siphonaptera and of 19 species of Diptera (see Supplementary Tables S1–S3). Specimen preservation, RNA extraction, cDNA library construction, DNA sequencing on Illumina HiSeq200/2500 and 4000 platforms, *de novo* assembly, and removal of possible contaminants was done as described by Misof et al. (2014), Peters et al. (2017), and Pauli et al. (2018). We combined our data with previously published nucleotide sequence assemblies of Antliophora transcriptomes (ten dipterans, three mecopterans, and three flea species; Misof et al., 2014; Peters et al., 2014; Pauli et al., 2018; Kutty et al., 2019) and with data from draft genomes genomes (four dipterans). In addition, we added transcriptome nucleotide sequence data from four lepidopteran and two trichopteran species (Misof et al., 2014) that we used as outgroups. Gene orthology was inferred using the ortholog set published by Pauli et al. (2018) comprising 3,145 nuclear single-copy protein-encoding genes (hereafter referred to as ortholog groups, OGs) and based on the official gene sets of three Diptera species and two species of Lepidoptera (Pauli et al., 2018).

We used Orthograph v. 0.5.4 (Petersen et al., 2017) to identify orthologous transcripts, and we only kept species from which we had at least 60% of the 3,145 single-copy nuclear genes of the ortholog set. This was the case for four fleas, twelve mecopterans, 32 dipterans, and eight outgroup species, see Supplementary Table S4. Multiple sequence alignment of OGs, alignment refinement, removal of outlier sequences (Supplementary Table S5), identification of protein domains, removal of ambiguously aligned sections, and the calculation of information content of partitions followed the procedures published by the 1KITE consortium (*e.g.*, Misof et al, 2014; Peters et al., 2017; Simon et al., 2018). Details are provided in the Supplememtary Materials, sections S1–S4.

After removal of data blocks (based on protein domains) with zero information content, we compiled three concatenated datasets for further analyses: i) all data blocks with an information content > 0 resulted in a dataset, [hereinafter called dataset] 0, with 56 species, 3,979 data blocks and a matrix length of 1,431,730 sites at the amino acid level. ii) a second dataset was compiled with the software MARE v. 0.1.2-rc (Misof et al., 2013) selecting an optimal subset (SOS), hereinafter called dataset SOS (53 taxa, 2,505 data blocks and 869,055 aligned amino acids). iii) with the third dataset, we followed the rationale of Dell’Ampio and colleagues (Dell’Ampio et al., 2014) and aimed to have a maximize overlap of taxa of interest. Therefore, we kept only data blocks that included all mecopteran species, all flea species, at least one representative of infraorders or superfamilies of included Diptera, and at least one representative of both outgroup taxa, Lepidoptera and Trichoptera (Supplementary Table S6). This resulted in our most optimized dataset, hereinafter called data set OPTI, including 56 species, 1,116 data blocks, spanning an alignment length of 683,836 amino acid sites. Supermatrices were analysed with MARE v. 0.1.2 -rc (Misof et al., 2013), AliStat v. 1.6 (Wong et al., 2020), and SymTest v. 2.0.44 (Jermiin & Ott, 2017) to assess overall information content, data coverage, and to explore whether our datasets were consistent with the assumption of historically stationary, time-reversible, and homogeneous (SRH) conditions (Ho & Jermiin, 2004; Jermiin et al., 2004; Ababneh et al., 2006). We restricted analyses at nucleotide level to our most optimized dataset OPTI. We only kept the 2^nd^ codon positions during tree inference and all subsequent analyses to reduce possible misleading effects due to saturation and/or violation of SRH conditions (see Supplementary Materials, section S5). An overview of analyzed datasets provided in Table 2.

**Table 2.** Overview of final datasets and analyses. Further dataset diagnostics are in detail described in the Methods Summary and in the Supplementary Materials, Supplementary Table 7. Note that the number of included partitions does not refer to single copy genes or protein domains: they represent the number of partitions after merging data blocks with PartitionFinder to find the optimal partition scheme. The overall coverage refers to data blocks. NA: not applicable.

| **Dataset** | **Data type** | **# Species** | **# Sites** | **# Data blocks** | **Overall coverage** | **# Partitions** | **Analyses** |
| --- | --- | --- | --- | --- | --- | --- | --- |
| dataset 0 | amino acid level | 56 | 1,431,730 | 3979 | 81.0% | 1,907 | ML tree inference |
| dataset SOS | amino acid level | 53 | 869,055 | 2505 | 91.6% | 1,203 | ML tree inference |
| dataset OPTI_aa | amino acid level | 56 | 683,836 | 1116 | 98.1% | 519 | ML and Bayesian tree inference, AU tests, FcLM, $\Delta pL$ |
| dataset OPTI_nt2 | nucleotide level (only 2<sup>nd</sup> codon positions) | 56 | 683,836 | NA (single block) | 98.1% | 179 | ML and Bayesian tree inference, AU tests, FcLM, $\Delta pL$ |

Model selection and data block merging into partitions was done with PartitionFinder v. 2.0.0, prerelease 11 (Lanfear et al., 2014; 2016). When analyzing the amino acid datasets, we restricted model selection to five amino acid substitution models (see Supplementary Materials, section S6) and the protein mixture model LG4X, which accounts for FreeRate heterogeneity (Le et al., 2012) applying the rcluster algorithm. When analyzing the most optimized nucleotide dataset OPTI, we applied the k-means algorithm by Frandsen et al. (2015) and the general time reversible (GTR) model.

Phylogenetic trees were calculated under the Maximum Likelihood optimality criterion using IQ-TREE (v. 1.4.2 and 1.4.4) (Nguyen et al., 2015; Chernomor et al., 2016) with a partition-based approach using the edge-proportional partition model to allow partitions to have evolved with different evolutionary rates (option -ssp). For the less stringent datasets 0 and SOS, we conducted 50 ML tree inferences with random start trees. For the most optimized dataset OPTI, at the amino acid level (hereinnafter called dataset OPTI_aa), we inferred 225 ML trees (75 with random start trees, 75 with a parsimony start tree, and 75 with a fixed start tree, assuming monophyletic Mecoptera). For the dataset OPTI at nucleotide level explicitly including 2^nd^ codon positions (hereinafter called dataset OPTI_nt2), we inferred 75 ML trees with random start trees. Statistical support was assessed from thorough non-parametric bootstrap replicates ensuring bootstrap convergence (Pattengale et al., 2010) as implemented in RaxML (Stamatakis 2014), v. 8.2.11. Statistical bootstrap support (BS) was mapped onto the ML tree with the best log-likelihood score respectively for each dataset. Additionally, we applied a single branch SH-like approximate likelihood ratio test (SH-aLRT) to obtain alternative support values for the best ML trees as described by Guindon et al. (2010). To test for rogue taxa, we applied RogueNaRok v. 1.0 (Aberer et al., 2013) with default settings providing the best inferred ML tree. In addition to the Maximum Likelihood-based approach, we inferred phylogenetic trees from our dataset OPTI_aa and dataset OPTI_nt2 (including only 2^nd^ codon positions) with a Bayesian approach using ExaBayes v. 1.5 (Aberer et al., 2014). We used a partition-based approach and three independent runs ensuring convergence for each dataset and subsequently calculated majority rule consensus trees with Bayesian posterior probability support. Further details on phylogenetic tree inference are provided in the Supplementary Materials, section S7.

To evaluate which of competing phylogenetic hypotheses is better supported, we applied two strategies focusing on the relationships among mecopteran species and addressing whether Mecoptera are monophyletic using our most optimized datasets OPTI_aa and OPTI_nt2: first, we applied the Approximate Unbiased (AU) tests (Shimodaira 2002) at amino acid and at nucleotide level as implemented in IQ-TREE v. 1.6.12 testing our best ML topologies. Secondly, we applied Four-cluster Likelihood Mapping (FcLM) (Strimmer & v. Haeseler, 1997; Misof et al., 2014) as implemented in IQ-TREE v. 1.4.4. Finally, to identify possible confounding signal, three FcLM permutation approaches were applied as introduced in previous phylotranscriptomic studies (*e.g.,* Misof et al., 2014; Peters et al., 2017; Simon et al., 2018) to assess the possible effects of among-lineage heterogeneity and/or non-randomly distributed missing data present in our datasets. In short, permutation scheme I removed existing phylogenetic signal, but left the sequence position of missing data as well as amino acid frequencies of all sequences/lineages untouched. Permutation scheme II was similar to permutation scheme I, but removed signal from amino acid frequencies in sequences and lineages (by replacing non-ambiguous amino acid residues with once drawn with frequencies given in the LG model) making the dataset homogeneous in terms of not violating SRH conditions. Permutation scheme III was similar to permutation scheme II, but additionally randomized the distribution of missing data. If any analysis of these “permuted” datasets would result in obvious signal for a particular hypothesis, it would indicate that such support was driven by bias in the original data (details see Supplemenatry Materials, section S8).

To evaluate whether specific partitions support mecopteran monophyly or paraphyly, we evaluated single partitions in the dataset OPTI_aa and dataset OPTI_nt2 as described by Shen et al. (2017) and Simon et al. (2018). We considered two contrasting topologies: either mecopteran paraphyly with Nannocoristidae as the closest relatives of fleas or monophyletic Mecoptera with Nannochoristidae as sister group to remaining mecopterans. We calculated the partition log-likelihood (pL) scores using IQ-TREE v. 1.4.4 (option -wpl) for the two different tree topologies. We then evaluated the contribution of each partition based on the partition log-likelihood scores by calculating the difference of each pL score, given the best and the alternative topology for each partition (ΔpL_i_), see details provided by Simon et al., 2018 and in the Supplementary Materials, section S8). ΔpL scores were summed up to explore the overall signal in the dataset OPTI_aa and in the dataset OPTI_nt2 (details see Supplemenatry Materials, section S8).

To consider the interpretation of morphological characters and the monophyly of Mecoptera, the parsimonious tree lengths of the two alternative topologies (ML tree of dataset OPTI_aa versus ML tree OPTI_nt2) were calculated with Mesquite (v. 3.51; Maddison & Maddison, 2018), with a morphological dataset of 246 characters. The dataset was composed of relevant characters of immature stages and adults published in Beutel et al. (2011), Schneeberg and Beutel, (2011) and Friedrich et al., (2013); details are providd in Supplememtary Materials, section S9.

Details on all steps of the analyses are provided along with results and statistics in the Supplementary Text, in the Supplementary Tables, and in the Supplementary Figures.

Transcriptome raw reads and assembled transcriptomes are available at NCBI through the respective accession numbers (see Table 1 and Supplementary Tables S1–S3) and under the Umbrella BioProject “The 1KITE project: evolution of insects”. Supplementary data are available as Supplementary Archives on the DRYAD digital repository available with this study.

## Results and Discussion

Our phylogenetic analyses show maximal support for a sister group relationship between Diptera and the remaining antliophoran lineages (Figure 2). This result was obtained irrespective of the specific datasets or of the type of data that we analyzed or of the analytical approach that we applied (spanning ~ 680,000 to 1.4 million sites, for details on the datasets see Supplementary Materials, Supplementary Table S7). The resolution of phylogenetic relationships within Mecoptera and the placement of Siphonaptera, however, remained ambigious. Our inferred phylogenetic relationships are inconsistent regarding the monophyly of Mecoptera and the placement of Boreidae, Nannochoristidae, and Siphonaptera, and the inferred phylogenetic relationships depended on whether the data is analyzed at the amino acid or at the nucleotide level and on what method of tree inference we applied. ML trees inferred from the amino acid datasets suggest mecopteran paraphyly (92% BS, dataset OPTI_aa or even higher BS support in datasets 0 and SOS), with the enigmatic Nannochoristidae placed as sister group to Siphonaptera (Figure 2a). However, ML analyses of dataset OPTI at nucleotide level, (*i.e.* dataset OPTI_nt2), as well as Bayesian analyses of the dataset OPTI_aa and the dataset OPTI_nt2 recover a monophyletic Mecoptera, with Nannochoristidae as sister to all remaining families, though with low or moderate support (nucleotide dataset: ML: 67% BS, Figure 2b, Bayesian: amino acid dataset: BPP 0.67; nucleotide dataset BPP 0.94, Supplementary Figures). Since Bayesian posterior probabilities are known to provide inflated support in phylogenomic analyses (Yang & Zhu, 2018), we chose to explore and further analyze only our ML results based on the datasets OPTI_aa and OPTI_nt2.

**Figure 2.**
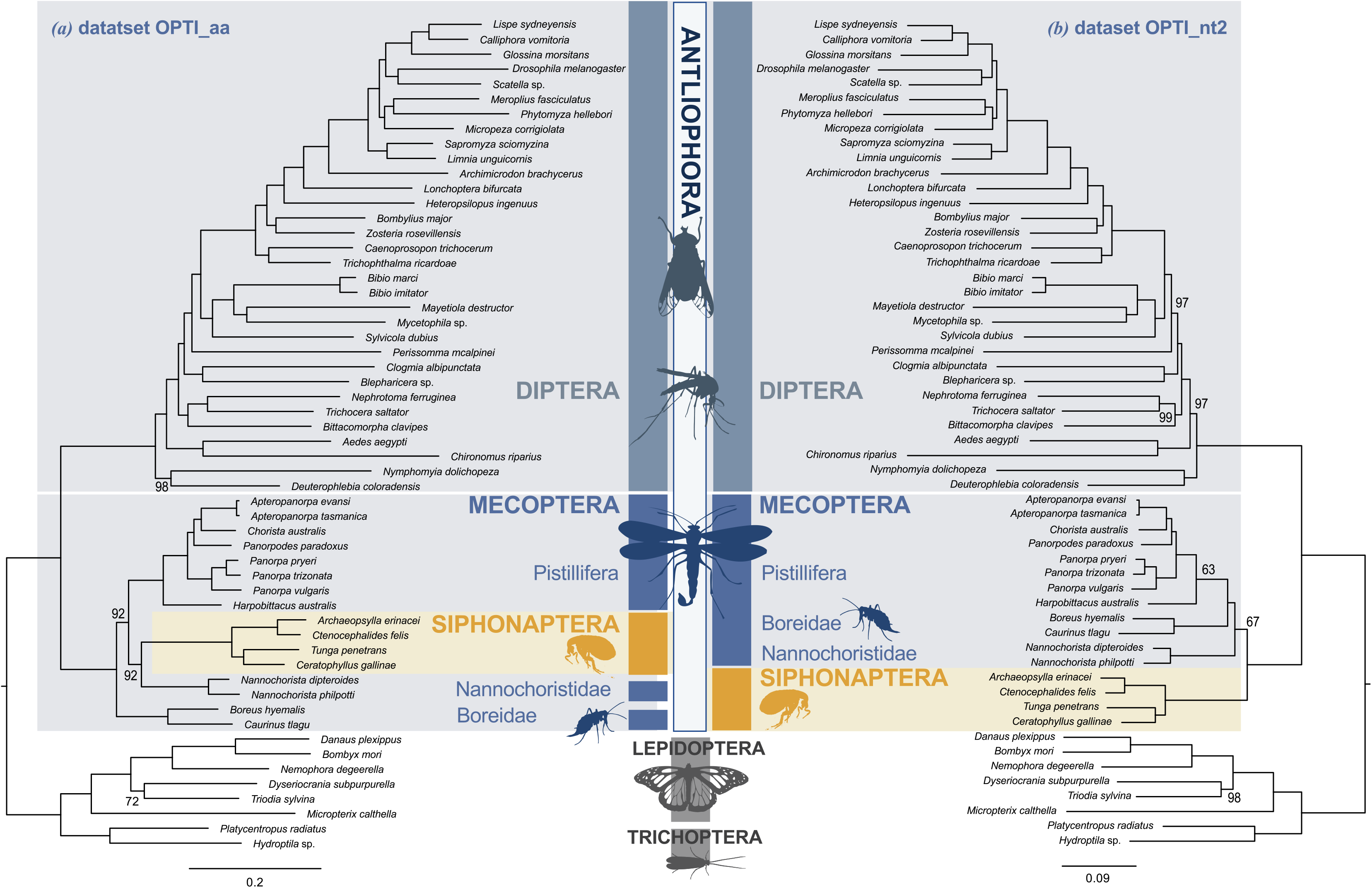
Contrasting maximum-likelihood phylogenies based on the most optimized datasets. Both Maximum-Likelihood (ML) trees were inferred with IQ-TREE and rooted with Lepidoptera + Trichoptera. Statistical support below 100% is displayed in numbers (%), all other splits are maximally supported. **(a)** Best Maximum-Likelihood (ML) tree inferred derived from our amino acid dataset OPTI_aa (56 taxa, alignment length: 683,836 amino acid sites, 519 partitions) with bootstrap support (BS) derived from 100 non-parametric bootstrap replicates. Mecopterans are paraphyletic with fleas nested inside and with Nannochoristidae as their closest relatives (Boridae (Pistillifera (Nannochoristidae, Siphonaptera))). **(b)** Best Maximum-Likelihood tree derived from our nucleotide dataset OPTI_nt2 (alignment length: 683,836 nucleotide site; 2^nd^ codon positions only, 179 partitions) with BS derived from 100 non-parametric bootstrap replicates. Mecoptera is monophyletic with Nannochoristidae being sister to a clade (Boridae, Pistillifera).

Our results reflect the complex history of conflicting hypotheses for mecopteran and siphonapteran phylogenetic relationships. Though previous studies based on molecular and morphological data have addressed the possibility of a paraphyletic Mecoptera that includes Siphonaptera, it was previously suggested that Boreidae are the sister group of Siphonaptera (Whiting 2002). Boreids and Siphonaptera are both small, flightless, jumping insects. However, structural affinities are vague and clearly defined synapomorphies are lacking (Beutel et al., 2008, 2009; Beutel & Friedrich, 2019). None of our analyses supported Boreidae as sister group to Siphonaptera. Instead, we found a weakly supported sister group relationship between fleas and the rare and phylogenetically ambiguously placed Nannochoristidae (Figures 1d and 2).

### Testing Competing Hypothesis and Putative Bias

#### AU Tests

We explored the signal in our data for different phylogenetic placements of Siphonaptera using multiple approaches. First, we tested for rogue taxa. Taxa that are unstable in their phylogenetic placement are known to reduce branch support (Aberer et al., 2013). We found that our analyses did not suffer from rogue taxon behavior. Next, we examined the effect of the starting tree on resulting topologies in our phylogenetic analyses in respect of dataset OPTI_aa. Irrespective of using either a random, a parsimony, or a fixed starting tree with monophyletic Mecoptera, all inferred ML trees from dataset OPTI_aa consistently yielded one unique topology with mecopteran paraphyly, and Nannochoristidae being the closest relatives of fleas (Figure 2a). To assess the magnitude of the signal for conflicting topologies in our datasets, we examined the 100 bootstrap trees derived from analyzing the amino acid dataset OPTI_aa and the dataset OPTI_nt2. Only 8% of the bootstrap trees inferred from dataset OPTI_aa provided support for monophyletic Mecoptera (counter to the best ML tree inferred from dataset OPTI_aa), while 36% of the bootstrap trees inferred from dataset OPTI_nt2 provided support for mecopteran paraphyly (counter to the best inferred from dataset OPTI_nt2). This indicates that signal is stronger for mecopteran paraphyly in dataset OPTI_aa than that for monophyletic Mecoptera in dataset OPTI nt2. However, AU tests comparing the log-likelihoods of two tree topologies offered no clarification. AU tests based on dataset OPTI_aa found that monophyletic Mecoptera as recovered in the best ML tree inferred from dataset OPTI_nt2 with Nannochoristidae sister to a clade comprising Boreidae and Pistillifera could not be rejected (p > 0.05). AU tests based on dataset OPTI_nt2 found that mecopteran paraphyly with Nannochoristidae as sister to fleas, as found in all ML trees inferred from dataset OPTI_aa (and from both less stringent datasets 0 and SOS) also could not be rejected (p > 0.05). Details are provided in the Supplementary Materials, section S8.

#### Four-cluster Likelihood Mapping: Alternative and Confounding Signal

To visualize differences in the distribution of signal for competing hypotheses of relationships between mecopteran lineages and Siphonaptera, we performed Four-cluster Likelihood Mapping (FcLM) on our original dataset OPTI_aa and OPTI_nt2. We additionally explored the presence of confounding signal (*i.e.*, non-randomly distributed data and or among-lineage heterogeneity) with three FcLM permutation approaches applied to each of the OPTI datasets that might affect our phylogenetic inference (*e.g.*, Misof et al., 2014; Peters et al., 2017; Simon et al., 2018). We defined four lineages: Nannochoristidae, Boreidae, Pistillifera, and Siphonaptera. This arrangement did not allow us to test for a monophyly of Mecoptera due to the unrooted nature of the resulting quartets (see also Szucsich et al., 2020 for a similar rooting issue in myriapod relationships). However, it enabled us to re-examine varying signal for the phylogenetic relationship between Nannochoristidae and Siphonaptera on one hand and that between Boreidae and Siphonaptera on the other (as hypothesized by Whiting 2002). Our FcLM analyses based on dataset OPTI_aa and dataset OPTI_nt2, both showed 100% quartet support for an assemblage of Nannochoristidae and Siphonaptera, and then for an assemblage of Boreidae and Pistillifera (Figure 3, upper quartet topology), respectively. Thus, we can clearly reject a close phylogenetic relationship of Boreidae and Siphonaptera (Figure 3, quartet topology 3 on the left).

**Figure 3.**
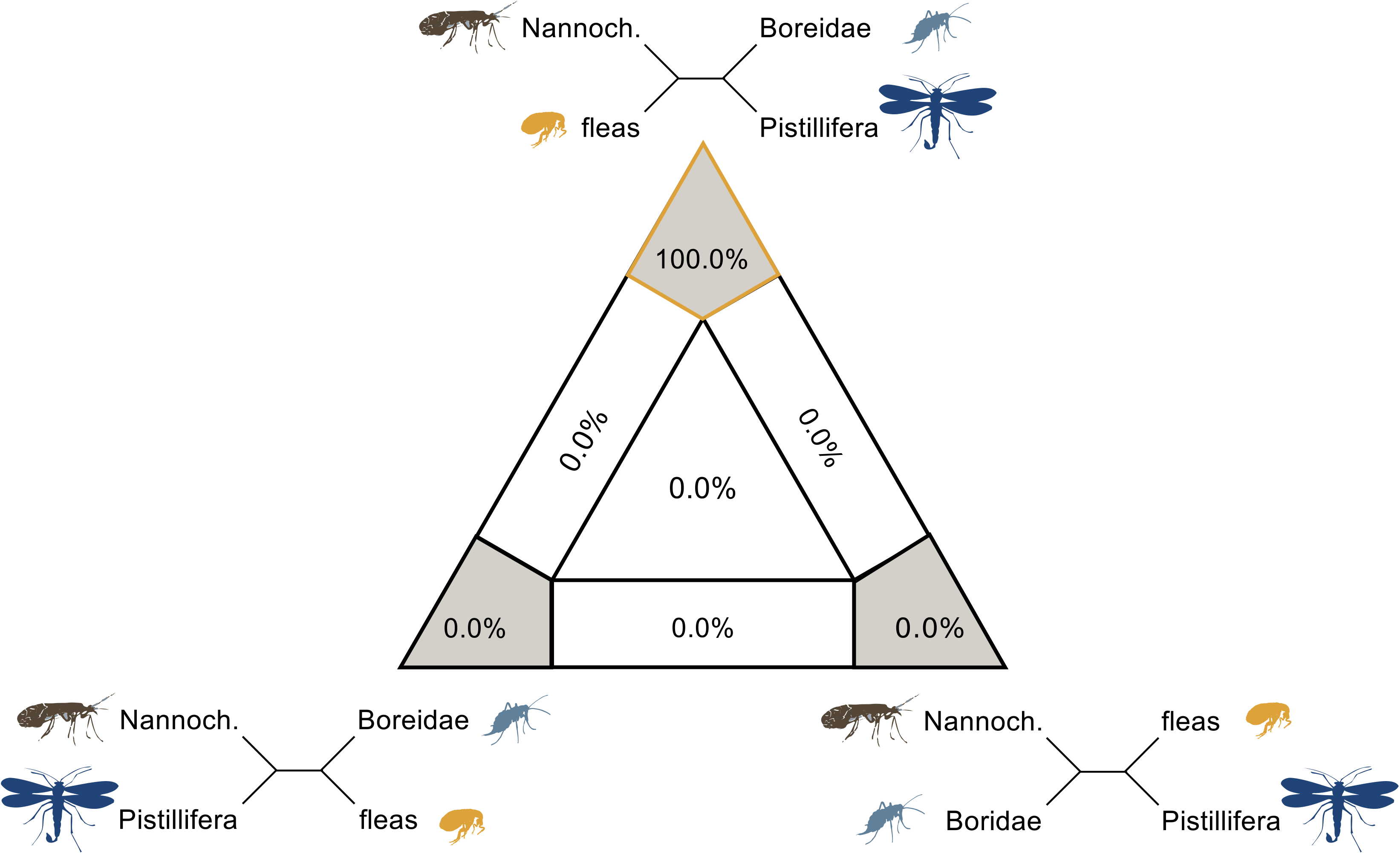
Four-cluster likelihood mapping of competing topologies for the placement of Siphonaptera. Quartet proportions (in %) are mapped on a 2D-simplex graph. We find maximal support for a close relationship between Siphonaptera and Nannochoristidae at amino acid and at nucleotide level (quartet topology on the top), rather than Boreidae (quartet topology on the left). Confounding signal could be excluded, see Supplementary Materials, section S8.

Results of the permutation approaches did not indicate confounding signal strongly affecting our original data analyses. Thus, we conclude that our datasets OPTI_aa and OPTI_nt2 are not biased by uneven data-distribution or by model violation due to among-lineage heterogeneity and consider our quartet mapping results reliable.

### Signal Considering Single Partitions: Partition Log-Likelihood Scores

Evaluating single partition log-likelihood (pL) scores for conflicting topological hypotheses allows for characterization of competing signal in phylogenomic datasets (Shen et al., 2017, Simon et al., 2018). For the dataset OPTI_aa, we evaluated 519 partitions previously used for tree inference. We compared the pL scores for two trees: the best ML tree with mecopteran paraphyly (Boreidae, ((Nannochoristidae, fleas), Pistillifera)) and the next most frequently recovered bootstrap tree with monophyletic Mecoptera (Nannochoristidae (Pistillifera, Boreidae)) as sister to fleas. By calculating and plotting the difference between the pL scores, (*i.e*., the ΔpL_i_, for each partition; Simon et al., 2018) comparing the two topologies, we found largely identical support for both trees (Figure 4a and Supplementary Materials, section S8 and Supplementary Table S8). The number of partitions and their cumulative length favoring mecopteran paraphyly was slightly smaller than the number of partitions and the cumulative length favoring monophyletic Mecoptera. However, the sum of ΔpL (see Simon et al., 2018) revealed a stronger signal for mecopteran paraphyly than for monophyletic Mecoptera. Except for three partitions strongly favoring mecopteran paraphyly, the signal favoring one topology over the other was in general quite low. Even less signal was present inspecting the dataset OPTI_nt2: we examined 179 partitions previously used for tree inference and found equivocal support for both the best ML tree (Figure 2b) inferred from dataset_OPTI_nt2 with monophyletic Mecoptera and best ML tree inferred from dataset OPTI_aa supporting the phylogenetic relationship (Boreidae ((Nannochoristidae, fleas) Pistillifera))) (Figure 2a). In fact, the largest partition, comprising 337,392 sites, does not favor one topology over the other (ΔpL=0; Figure 4b, Supplementary Table S9). There was only one partition spanning ~ 4,000 bp that contained very strong signal for monophyletic Mecoptera and only two partitions (spanning ~ 2,200 bp) that contained very strong signal for mecopterans being paraphyletic. However, summing up the ΔpL over all partitions, the signal was stronger for monophyletic Mecoptera. We found, however, no pattern favoring either para- or monophyletic mecopterans in terms of partition length, information content and/or partition-specific rates pointing to fast- or slow-evolving partitions. This suggests that ambiguity surrounding mecopteran monophyly is not driven by only a few partitions (see Shen et al., 2017), but instead is found throughout our transcriptome data, though overall signal is quite low.

**Figure. 4.**
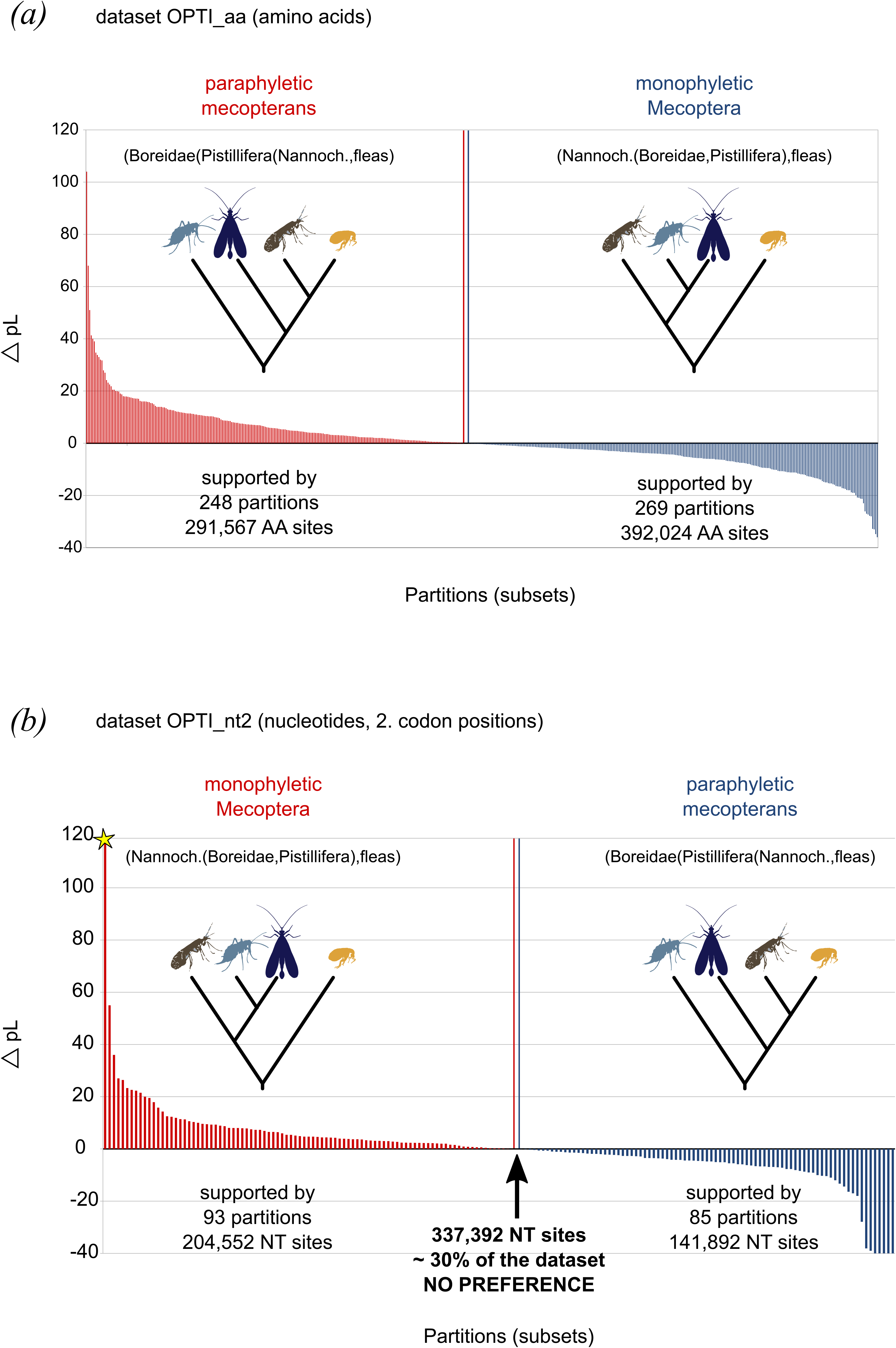
Difference of partition log-likelihood scores (ΔpL) for mecopteran paraphyly versus monophyly. Representation of the difference of each partition log-likelihood score (ΔpL_i_) received for either the topology suggesting mecopteran paraphyly or monophyly. The x axes show the partitions with the respective ΔpL_i_ displayed on the y-axes. Bars in red (positive scores) always show the support for the best ML tree inferred from the respective dataset (mecopteran paraphyly in dataset OPTI_aa vs. monophyletic Mecoptera in dataset OPTI_nt2). Bars in blue (negative scores) show support for the alternative topology (monophyletic Mecoptera in dataset OPTI_aa vs. mecopteran paraphyly in dataset OPTI_nt2) which could not be rejected by the AU test. **(a)** Partition log-likelihood scores (ΔpL) inferred from dataset OPTI_aa; summing up all ΔpL indicate a much stronger signal for paraphyletic Mecoptera (details see Supplementary Materials, section S8 and Table S8. **(b)** Partition log-likelihood scores (ΔpL) inferred from the dataset OPTI_nt2 (2^nd^ codon positions only); summing up all ΔpL indicate a stronger signal for monophyletic Mecoptera. Note that the partition with the strongest signal (4,755 units, length: 4,124 sites) in favor of monophyletic Mecoptera is indicated by a star since its value exceeded the scale shown here. Approximately half of the dataset (Subset #4 with a partition length of > 337,000 sites had a ΔpL = 0) showed no preference (details see Supplementary Materials, section S8 and Supplementary Table S9).

#### Morphological Interpretations

The monophyly of a clade comprising Siphonaptera and Mecoptera is unambiguously supported by all datasets analyzed irrespective of the data type or the applied methodological approach. This is in contrast to the morphology-based study by Beutel et al. (2011), in which fleas were found as sister to Diptera. This affinity to Diptera was mainly based on the absence of larval legs and adaptations of the adult mouthparts for liquid feeding, in the latter case implying parallel evolution in Nannochoristidae (Beutel & Baum, 2008). Morphological synapomorphies for a clade comprising Siphonaptera and Mecoptera are not convincing. The presence of an intraprofurcal muscle (Friedrich & Beutel, 2010) and of acanthae in the proventriculus (Richards & Richards, 1969) are non-reductive characters supporting this clade. However, Nannochoristidae lack the latter cuticular projections, presumably by secondary reduction. Another derived feature occurring in both Siphonaptera and Mecoptera is the loss of the outer circle of nine microtubules in the sperm axoneme (Dallai et al., 2003), except for Bittacidae in which this circle is present. The condition in Nannochoristidae is still unknown (Kristensen 1999; Gottardo et al., 2016). Using the extensive morphological dataset published by Beutel et al. (2011), the scenario with monophyletic Mecoptera (Figure 2b, right topology) requires ten fewer steps, compared to paraphyletic Mecoptera with fleas as subordinate group (Figure 2a, left topology). With a dataset focused on Mecoptera (Friedrich et al., 2013), monophyletic Mecoptera requires nine steps less than a paraphyletic Mecoptera scenario. Even though these results seem to favor monophyletic Mecoptera quite clearly, there is some morphological evidence for the evolution of Siphonaptera from within Mecoptera. We have summarized the morphological evidence for both monophyletic (Scenario 1) and paraphyletic (Scenario 2) mecopterans in Table 3. Further details can be found in the Supplementary Materials, section S9 and Supplementary Tables S10–S11.

**Table 3.** Summary of morphological implications of Mecoptera scenarios.

| **Character** | **Scenario 1**<br>**Paraphyletic “Mecoptera”**<br>**(incl. fleas)** | **Scenario 2**<br>**Monophyletic Mecoptera and**<br>**basal Nannochoristidae** |
| --- | --- | --- |
| Labral and lacinial food channels | synapomorphy of Nannochoristidae and Siphonaptera [convergently present in some Diptera] | parallel evolution in Nannochoristidae and Siphonaptera |
| Secretion former muscle | independently developed in Pistillifera and Boreidae | synapomorphy of Pistillifera and Boreidae |
| Thickening of mandibular tendon (“Apodemwalze”) | parallel evolution in Pistillifera and Boreidae | synapomorphy of Pistillifera and Boreidae |
| Elongated rostrum | independently developed in Pistillifera and Boreidae | synapomorphy of Pistillifera and Boreidae [reversal in *Caurinus*] |
| Proventriculus with acanthae | synapomorphy of Mecoptera and Siphonaptera (reduction in Nannochoristidae) | synapomorphy of Mecoptera and Siphonaptera (reduction in Nannochoristidae) |
| Postcerebral pharyngeal pumping chamber | synapomorphy of Nannochoristidae and Siphonaptera [convergently present in some Diptera] | parallel evolution in Nannochoristidae and Siphonaptera (or secondarily lost in Boreidae + Pistillifera) |
| Intraprofurcal muscle | synapomorphy of “Mecoptera” and Siphonaptera | synapomorphy of Mecoptera and Siphonaptera |
| Specific sperm pump (Mickoleit 2009) | synapomorphy of Nannochoristidae + Siphonaptera | independent evolution in Nannochoristidae and Siphonaptera (or secondarily lost in Boreidae) |
| Larvae with orthognathous head, grub-like body with folds and protuberances | parallel evolution in Pistillifera and Boreidae | synapomorphy of Pistillifera and Boreidae |
| Larval compound eyes | synapomorphy of Pistillifera + (Nannochoristidae + Siphonaptera) [with loss of eyes in Siphonaptera] | independent evolution in Pistillifera and Nannochoristidae (or reversal to stemmata in Boreidae) |
| Larval legs absent | independent loss in Diptera and Siphonaptera | independent loss in Diptera and Siphonaptera |

#### Morphological Characters Supporting Monophyletic Mecoptera

The unique “Sekretformer” muscle, an intrinsic muscle of the salivarium in the adult, appears to be a convincing synapomorphy of a clade (Boreidae+Pistillifera), with a clearly plesiomorphic condition in Nannochoristidae, with the ancestral set of three muscles of the apical salivary duct (Beutel & Baum, 2008). Other potential apomorphic larval characters supporting Mecoptera– excluding Nannochoristidae–are for instance an orthognathous head, and a grub-like unsclerotized postcephalic body with folds and protuberances (Fabian et al., 2015). Pistillifera have secondary larval compound eyes, and the vestigial lateral eyes of nannochoristid larvae were also interpreted as reduced compound eyes by Melzer et al. (1994). Typical holometabolan stemmata are present in larvae of Boreidae (Fabian et al., 2015), while larval eyes have been lost in Siphonaptera.

#### Morphological Synapomorphies Supporting Paraphyletic Mecoptera

Our ML trees based on amino acid data place Siphonaptera within Mecoptera in a sister-group relationship with Nannochoristidae. Some potential synapomorphies between Nannochoristidae and Siphonaptera have been proposed, from the adult male sperm pump, the adult head, and the larvae: the sperm pumps of Siphonaptera and Nannochoristidae have multiple similarities. Potential synapomorphies are a pistil chamber moved against a fixed pistil, the formation of a strongly sclerotized fulcrum supporting the roof of the endophallus, and the formation of a fixed pistil by the endophallic roof and the fulcrum (Mickoleit 2009; Beutel & Friedrich, 2019). Fleas and nannochoristids also share several features of the immature stages. Their larvae are very slender, with a prognathous head and a cylindrical body with a very smooth cuticular surface (Beutel & Friedrich, 2019) which is in striking contrast to the larvae of Boreidae and Pistillifera (*e.g.*, Byers 1987; Fabian et al., 2015; Beutel & Friedrich, 2019). However, a prognathous head is very likely part of the groundplan of Antliophora and Mecopterida (*i.e.*,Antliophora, Lepidoptera and Trichoptera, Beutel et al., 2011), and a slender and smooth postcephalic larval body is also common among dipteran groups. Consequently, these features are most likely plesiomorphic.

In summary, quantitative analysis of morphological characters supports the monophyly of Mecoptera more than their paraphyly. However, there are some characters that could be interpreted as supporting mecopteran paraphyly, with subordinate Siphonaptera. Both the monophyletic and paraphyletic Mecoptera scenarios imply some independent evolution and/or loss of complex morphological characters. For example, if Mecoptera are monophyletic then a similar sperm pump must have evolved independently in Nannochoristidae and fleas, or it has been lost in Boreidae. If Mecoptera are paraphyletic then the “Sekretformer” muscle of the salivarium has evolved independently in Boreidae and Pistillifera. Like the molecular characters, morphological characters do not provide us with compelling evidence to decide whether Mecoptera are monophyletic or paraphyletic.

### Evolutionary History of Mecoptera and Siphonaptera

Although our data do not resolve the question of mecopteran monophyly, it settles on two of 32 possible resolutions of this 5-taxon problem comprising the lineages flies, fleas, Nannochoristidae, Boreidae and the remaining Mecoptera families, *i.e.* Pistillifera. Our results show that fleas arose either before all families of crown Mecoptera, or as an early split among crown mecopteran families. The fossil record is consistent with either result, with the diversity of fossil mecopterans existing from the Permian through the Cretaceous, well-preserved fossil Boreidae from the Late Jurassic (*Palaeoboreus*), Nannochoristidae from the Early Jurassic, and unambiguous flea fossils from the Early Cretaceous (*Tarwinia,* Jell & Duncan, 1986; Huang 2015). Divergence time analyses date the split between Mecoptera and Siphonaptera during the Triassic and Early Jurassic (Misof et al., 2014). The evolution of flea hosts is also consistent with this timing (Zhu et al., 2015); extant fleas are mostly mammal parasites, lineages that feed on birds being derived from primarily mammal feeding ancestors (Zhu et al., 2015). Early mammals appeared in the Late Triassic, with the earliest placental mammals appearing in the Early Cretaceous (dos Reis et al., 2015). Whatever the precise phylogenetic topology may be, the emergence of Siphonaptera as a separate lineage is bounded by the Permian and the Early Jurassic.

Mecoptera are often considered a relict lineage whose peak of species richness was in the ancient past; yet, if fleas are truly a mecopteran lineage, then parasitism posed a new avenue for evolutionary success. The ecological specialization of fleas as mammalian ectoparasites likely had major impacts on the flea lineage: accelerated anatomical divergence from their relatives — including the loss of wings and eyes, extreme lateral compression, a well-developed jumping ability — and an expansion of species richness due to the availability of novel host niches. While Mecoptera include only approximately 600 described extant species, there are more than 2,000 described flea species. Though Mecoptera are an order with relatively few extant species, their fossil record shows three times as much diversity historically (Grimaldi & Engel, 2005). Nannochoristids had a previously broad geographic distribution that through extinction is now limited to South America and Oceania (Penny 1975). It is possible that the seemingly high incidence of extinction of mecopteran lineages, particularly those near Nannochoristidae, may have contributed to the ambiguity in discerning the phylogenetic relationships of Siphonaptera.

## Conclusions

As the field of molecular systematics has become increasingly data-rich and analytically sophisticated over the past decade, it seemed possible that large-scale genomic datasets from new sequencing technologies could resolve many if not all outstanding phylogenetic questions. Yet, for some regions of the insect tree of life fundamental questions remain about the relationships between orders, despite extensive phylogenomic analyses. This is the case in Antliophora. The extreme morphological divergence of Siphonaptera has obscured their true origins among the Antliophora, with almost every possible resolution of the Diptera, Siphonaptera, and Mecoptera trichotomy proposed by morphological studies (Whiting 2002), including the division of extant Mecoptera into three order-level lineages. Our analyses of phylogenomic data suggest that there is little signal even in very large datasets to unambiguously reconstruct the early branching of extant mecopteran lineages and Siphonaptera. Future work to resolve the question of mecopteran monophyly will likely need to rely on other character systems such as, for example, genomic meta-characters (see Niehuis et al., 2012) to arrive at a robust answer.

## Author’s Contributions

KM, MT, BMi, BMW and DKY conceived the study. KM, BMW, BMi, RGB, DKY, SG, KMB, RSP collected or provided samples. XZ and SL were responsible for sequencing and assembly. AD, LP and KM managed, processed and submitted the transcriptome data. Orthology assignment was done by KM, MP and ON. AD, LP, KM, DB, MP, CM, ON, RSP, and BMi contributed to the 1KITE pipeline development. KM did all molecular data analyses with the help of MT; OH helped with all HPC Cluster-related analyses and BQM. helped with phylogenetic analyses by debugging IQ-TREE. AK and BMo performed Bayesian tree inference. FF and RGB sampled and analyzed morphological data and provided interpretations. All authors contributed to the writing of the manuscript, with MT, KM BMW and DKY taking the lead.

## Acknowledgements

Data presented here are the result of the collaborative efforts of the 1KITE consortium (https://www.1kite.org/). The authors acknowledge all who helped to make this project happen and assisted with any aspects. Specifically, we thank Lars S. Jermiin, Michael Ott, Thomas Wong, Robert Lanfear, Brett Calcott, Guanliang Meng, and Hermes Escalona for help and discussion of with bioinformatic analyses; and we thank Robert Waterhouse and Alexandros Vasilikopoulos for providing the ortholog set. Special thanks for providing, help to collect or organize crucial species samples go to Phil Suter, Alexander Blanke, Julia Schwarzer and her toe, Greg W. Courtney, Urs Schmidt-Ott, David J. Ferguson, Matthew A. Bertone, Susanne Dobler, Derek S. Sikes, Loren K. Russell, Yuta Mashimo, Ryuichiro Machida, and Peter McQuillan. KM and DKY thank the Schlinger Foundation for funding a postdoctoral position. KM thanks her cat for all patience. Last but not least, we thank Hans Pohl for providing pictographs.

